# Development of a Coronavirus Disease 2019 Nonhuman Primate Model Using Airborne Exposure

**DOI:** 10.1101/2020.06.26.174128

**Authors:** Sara C. Johnston, Alexandra Jay, Jo Lynne Raymond, Franco Rossi, Xiankun Zeng, Jennifer Scruggs, David Dyer, Ondraya Frick, Joshua Moore, Kerry Berrier, Heather Esham, Joshua Shamblin, Willie Sifford, Jimmy Fiallos, Leslie Klosterman, Stephen Stevens, Lauren White, Philip Bowling, Terrence Garcia, Christopher Jensen, Jeanean Ghering, David Nyakiti, Stephanie Bellanca, Brian Kearney, Wendy Giles, Nazira Alli, Fabian Paz, Kristen Akers, Denise Danner, James Barth, Joshua A. Johnson, Matthew Durant, Ruth Kim, Margaret LM Pitt, Aysegul Nalca

## Abstract

Airborne transmission is predicted to be a prevalent route of human exposure with SARS-CoV-2. Aside from African green monkeys, nonhuman primate models that replicate airborne transmission of SARS-CoV-2 have not been investigated. A comprehensive and comparative evaluation of COVID-19 in African green monkeys, rhesus macaques, and cynomolgus macaques following airborne exposure to SARS-CoV-2 was performed to define parameters critical to disease progression and the extent to which they correlate with human COVID-19. Respiratory abnormalities and viral shedding were noted for all animals, indicating successful infection. Cynomolgus macaques developed fever, and thrombocytopenia was measured for African green monkeys and rhesus macaques. Type II pneumocyte hyperplasia and alveolar fibrosis were more frequently observed in lung tissue from cynomolgus macaques and African green monkeys. The data indicate that, in addition to African green monkeys, macaques can be successfully infected by airborne SARS-CoV-2, providing viable macaque natural transmission models for medical countermeasure evaluation.

**One Sentence Summary:** Nonhuman primates develop COVID-19 following airborne virus exposure.

## Main Text

Severe acute respiratory syndrome coronavirus 2 (SARS-CoV-2) is responsible for an ongoing worldwide pandemic of mild to severe respiratory disease identified as coronavirus disease 2019 (COVID-19). Clinical signs for symptomatic patients have ranged from a mild disease characterized by fever, cough, and/or increased respiratory rate to severe (and sometimes fatal) disease characterized by severe pneumonia, respiratory failure, sepsis/septic shock, and heart failure (*1–4*).

Early reports on transmission of SARS-CoV-2 suggested that human spread occurred primarily through respiratory droplets (particles created by coughing and sneezing) and direct contact (*2, 5–9*). However, recent evidence indicates that airborne transmission may actually represent the most prevalent human infection route (*10*). This route relies on virus-laden particles that are less than 5μm in size, which can travel extended distances in the air. To test and evaluate novel medical countermeasures against SARS-CoV-2, animal models that mimic a natural route of infection and recapitulate human disease are required. To date, countermeasure testing in nonhuman primates has predominantly utilized combination models, wherein high doses of virus in excess of 10^6^ TCID_50_ are inoculated intratracheal (IT)/intranasal (IN) or IT/IN/ocular/oral (*11–13*). Although animals in these experiments develop some mild clinical signs of respiratory disease, the routes of exposure are largely artificial in that they rely on direct contact inoculation rather than natural transmission. Herein, we sought to assess nonhuman primate airborne challenge models of SARS-CoV-2. Airborne virus was generated using a Collison nebulizer to produce a highly respirable target aerosol of 1 to 3μm mass median aerodynamic diameter. Healthy, SARS-CoV-2 serologically-naïve African green monkeys (AGM, n=3), rhesus macaques (RM, n=4), and cynomolgus macaques (CM, n=4) were exposed to SARS-CoV-2 on Study Day 1 by the aerosol route. The mean virus inhaled dose was 3.84×10^4^ plaque forming units (pfu), with AGM receiving an average of 3.80×10^4^ pfu, RM receiving an average of 2.87×10^4^ pfu, and CM receiving an average of 4.86×10^4^ pfu. Animals were observed daily for clinical signs of illness. All animals survived until the end of study (Study Day 18 or Study Day 19) at which time they were humanely euthanized.

Clinical observation parameters are summarized in Table 1. In general, clinical disease findings were noted as early as Study Day 2 and as late as Study Day 18. The earliest clinical sign was the development of fever (defined as a reading that was ≥1.5°C above baseline for a duration of greater than 2 hours by telemetry), which was consistently noted for all CM between Study Days 2-3 (Figure 1). The maximum temperature change for CM was between 2.2-3.0°C (mean = 2.5°C). Although small transient temperature increases were noted for some AGM and RM, none met study-defined criteria (≥1.5°C above baseline for a duration of greater than 2 hours) to be called fever. Therefore, fever was a finding exclusive to CM infected with airborne SARS-CoV-2.

**Table 1.** Summary of Clinical Disease Findings.

| Animal Number | AGM 1 | AGM 2 | AGM 3 | RM 1 | RM 2 | RM 3 | RM 4 | CM 1 | CM 2 | CM 3 | CM 4 |
| --- | --- | --- | --- | --- | --- | --- | --- | --- | --- | --- | --- |
| Elevated Temp +/- Fever |  |  |  |  |  |  |  | X | X | X | X |
| Mild Hypoxia<sup>1</sup> | X | X |  | X | X | X |  |  | X |  |  |
| Tachycardia |  | X |  | X |  |  |  | X | X |  | X |
| Increased Respiratory Rate |  |  |  |  |  |  |  | X | X |  |  |
| Increased Respiratory Sounds (Auscultation) | X | X | X | X | X | X | X | X | X | X | X |
| Cough | X |  |  |  |  |  |  |  |  |  |  |
| Lung Infiltrates |  | X | X | X |  |  | X | X | X | X | X |
| Lung Opacity | X | X | X | X | X | X | X | X | X | X | X |
| Evidence of Increased Respiratory Effort<sup>2</sup> | X | X | X |  | X |  |  | X | X |  |  |
| Clotting Abnormality |  |  |  |  |  |  |  |  |  |  |  |
| Unusual Bruising | X |  |  |  |  |  |  |  |  |  |  |
| Petechial Rash | X |  |  |  |  |  |  |  |  |  |  |
| Erythema of Eyes | X |  |  | X |  | X | X |  |  |  |  |
| Ocular Discharge | X |  |  |  |  |  |  |  |  |  |  |
| Lymphadenopathy |  |  | X |  | X |  |  |  |  |  | X |
| Rectal Bleeding |  |  |  |  | X | X | X |  | X | X |  |
| GI Gas/Fluid |  |  |  |  |  |  | X |  |  |  | X |
| Shaking |  |  |  |  |  |  |  |  | X |  |  |

<sup>1</sup>Mild hypoxia is defined as 90-94% SpO<sub>2</sub>
<sup>2</sup>Evidence of increased respiratory effort = abdominal component to breathing and/or nasal flare

**Fig. 1.**
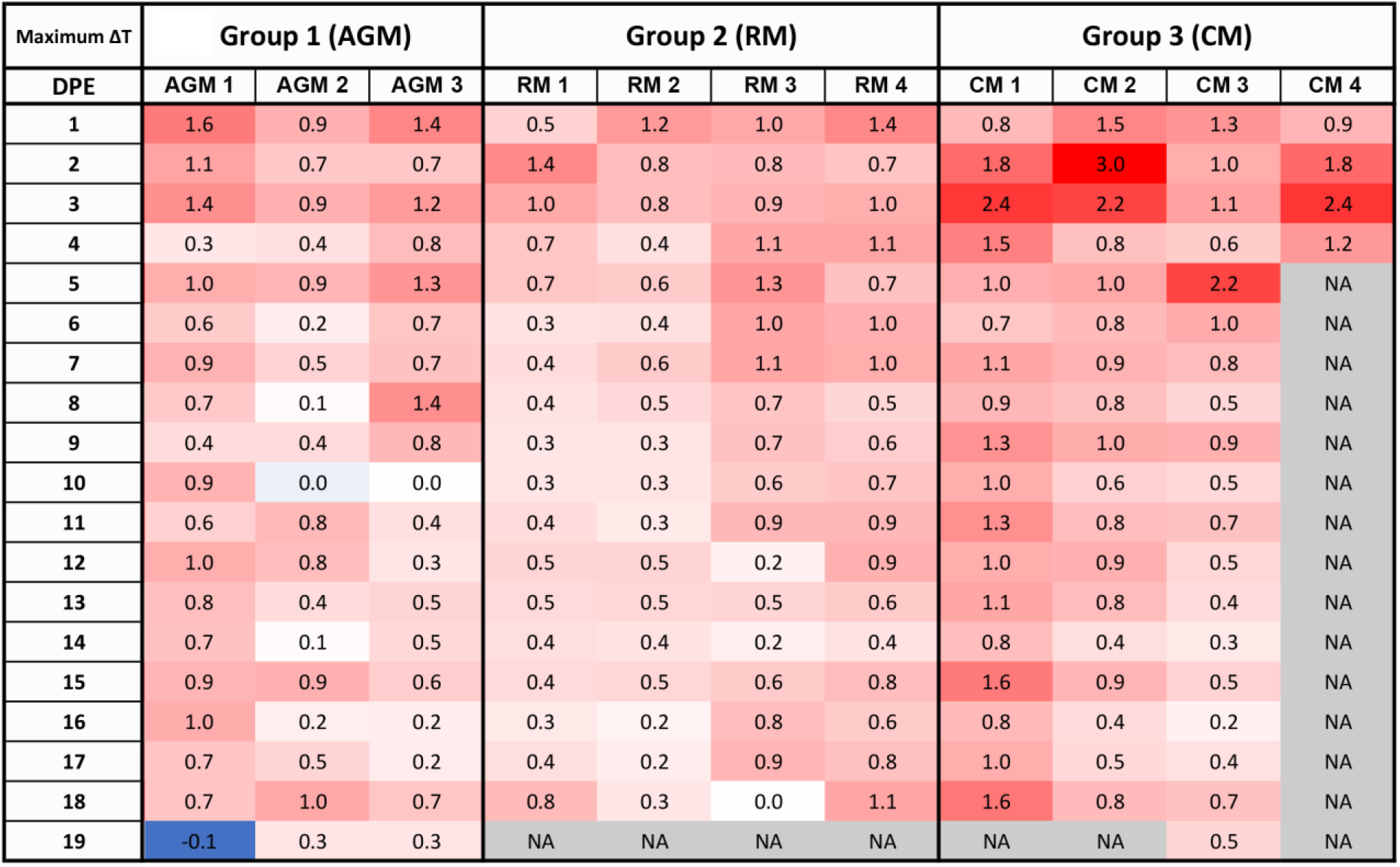
Fever responses as measured by telemetry. DSI M00 telemetry devices were used to collect body temperature data. The figure shows the maximum change in temperature (°C) for the 24 hour daily time period.

Other common clinical findings for animals on study included increased respiratory sounds on auscultation, and radiographic findings of increased lung opacity with or without the presence of infiltrates (Figure 2). Lung opacity was noted for all animals on study; however, lung infiltrates were more commonly noted for AGM and CM. Generally, radiographic findings were noted prior to audible lung sounds, with radiographic findings commonly identified between Study Days 3-11, and increased respiratory sounds noted between Study Days 5-9.

**Fig. 2.**
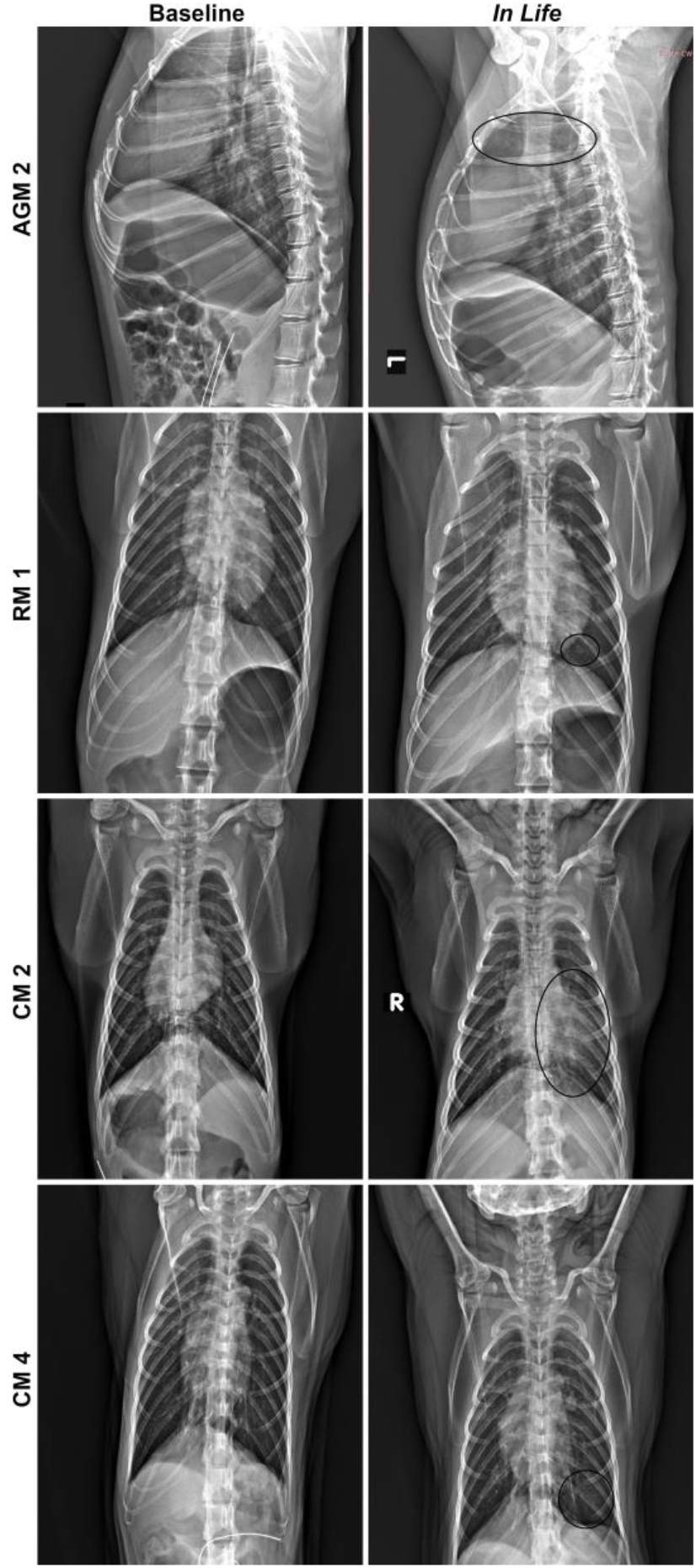
Radiographic changes for SARS-CoV-2 infected NHP. Left panels show baseline images collected prior to telemetry surgery, and right panels show images collected on Study Day 7. All images are ventrodorsal except for AGM 2 which are lateral images. AGM 2 = Infiltrates present in cranial lung lobes (circled). RM 1 = Infiltrate present in caudal left lung (circled). CM 2 = Infiltrate present in left lung (circled). CM 4 = Increased opacity in caudal left lung (circled).

For AGM, additional findings identified as consistent indicators of disease for this model included mild hypoxia and evidence of increased respiratory effort. For RM, additional consistent findings included mild hypoxia and erythema around the eyes. Rectal bleeding, which was a common finding for this group, was only noted on palpation and likely the result of repeated swabbing leading to irritation and mild bleeding. For CM, an additional consistent finding included tachycardia.

Disease severity based on clinical signs was graded using a scoring system that can be found in Table S1 [Note: this table was based on a table utilized by Finch *et al* (*14*) and information contained within (*15*) with certain updates based on the COVID-19 disease observed in the primate species being evaluated in the present study]. Average disease severity scores were as follows (Figure S1): AGM was 11.00 (range: 8-16), RM was 9.75 (range: 8-11), and CM was 14.00 (range: 10-19). The highest severity score on study was 19 (CM 2), and the lowest severity score on study was 8 (AGM 3 and RM 4).

Complete blood counts and clinical chemistries (Figure 3 and Table S2) were performed on whole blood and serum, respectively. Peak change from baseline (defined as data from Study Day 1) is represented, and Table S2 depicts the parameters for which changes from baseline were greater than or equal to 25%. Lymphocyte and neutrophil fluctuations were noted for most animals on study and were likely a result of the inflammatory response to infection.

**Fig. 3.**
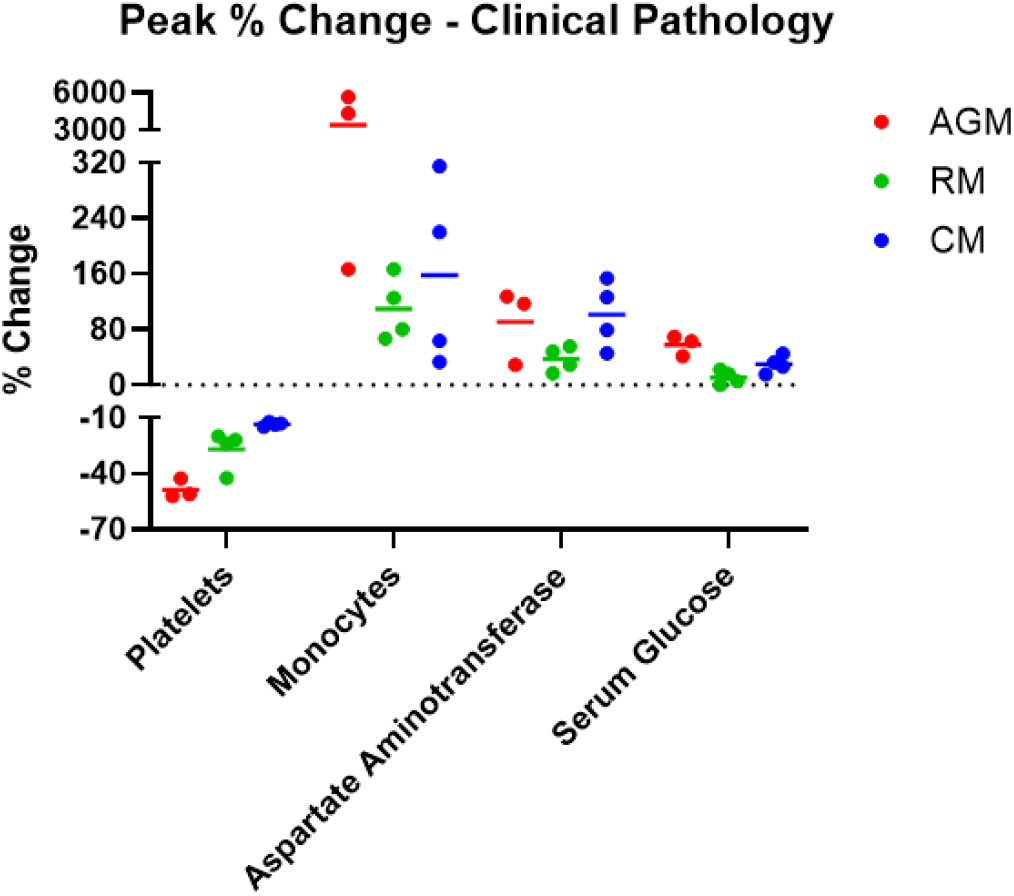
Hematology and Clinical Chemistries. Hematology was performed on EDTA whole blood using a HM5 instrument. Clinical chemistries were performed on serum using a Piccolo Point-Of-Care instrument. Measurements for animals on study are shown as percent change from baseline (Study Day 1 values for that animal) for peak values for each analyte.

AGM and RM exhibited a decrease in platelet counts beginning on Study Days 3-5 that persisted through Study Days 15-18. Nadirs were reached earlier in AGM (Study Day 5) and were more pronounced (40%) compared to RM. Decreases may relate to lung injury and subsequent sequestration; evidence of inflammation and chronic lesions noted histologically further support this assessment.

Increases in one or more liver-related enzyme activities were present for most animals, with the most notable and consistent changes occurring for AGMs and CMs. For CMs, elevated levels appeared to correlate with the timing of the fever response, with levels approximating baseline by Study Day 7. A more protracted response was noted for AGMs, with elevated levels noted between Study Days 5-11. Unfortunately, necropsy occurred 7-11 days after enzyme activities had returned to baseline, so any corresponding histologic changes in the liver that might have suggested a cause for the elevations were no longer present.

Increases in glucose concentration were noted in AGM over Study Days 3-7 (1.3 fold) and 15-18 (1.6 fold). A similar trend was noted for CMs, but the degree and consistency of the change was less pronounced. These findings may relate to a corticosteroid or epinephrine response. Changes in glucose levels were not noted for RMs.

Significant increases in monocytes, measured on Study Day 5, were present only for AGM; a 57 fold and 44 fold increase from baseline was noted for AGM 2 and AGM 3, respectively. Although the physiological relevance of this finding is unclear at this time, monocytosis was a consistent and exclusive finding for AGM. No other notable changes in clinical chemistry parameters were measured on this study.

Quantitative reverse transcription polymerase chain reaction (qRT-PCR) was performed on oropharyngeal (OP), nasopharyngeal (NP), and rectal swabs to assess viral shedding (Figure 4), and on whole blood to assess circulating viral load (data not shown). Viral RNA in most whole blood samples was not detected; when detected, it was on Study Day 11, was typically below the limit of quantification, and was most often noted for AGM. Viral RNA in rectal swabs, when noted, was measured on or after Study Day 7; this finding was most often noted for AGM. Viral RNA was detected for most animals by Study Day 3 in OP and NP swabs. Peak levels in NP swabs were between 9.63-11.02 Log_10_ genomic equivalents (ge)/mL, and peak levels in OP swabs were between 7.50-10.70 Log_10_ ge/mL. Significant differences in peak levels between swab types and groups were not observed with one exception: CM. For this group, peak levels in OP swabs were 1-2 logs lower compared to OP swab peak levels for other groups. For the majority of the animals, viral RNA was no longer detected by Study Day 11. Viral RNA present for a few AGM on Study Day 18 likely represented residual RNA in the nasal cavity and not actively replicating virus.

**Fig. 4.**
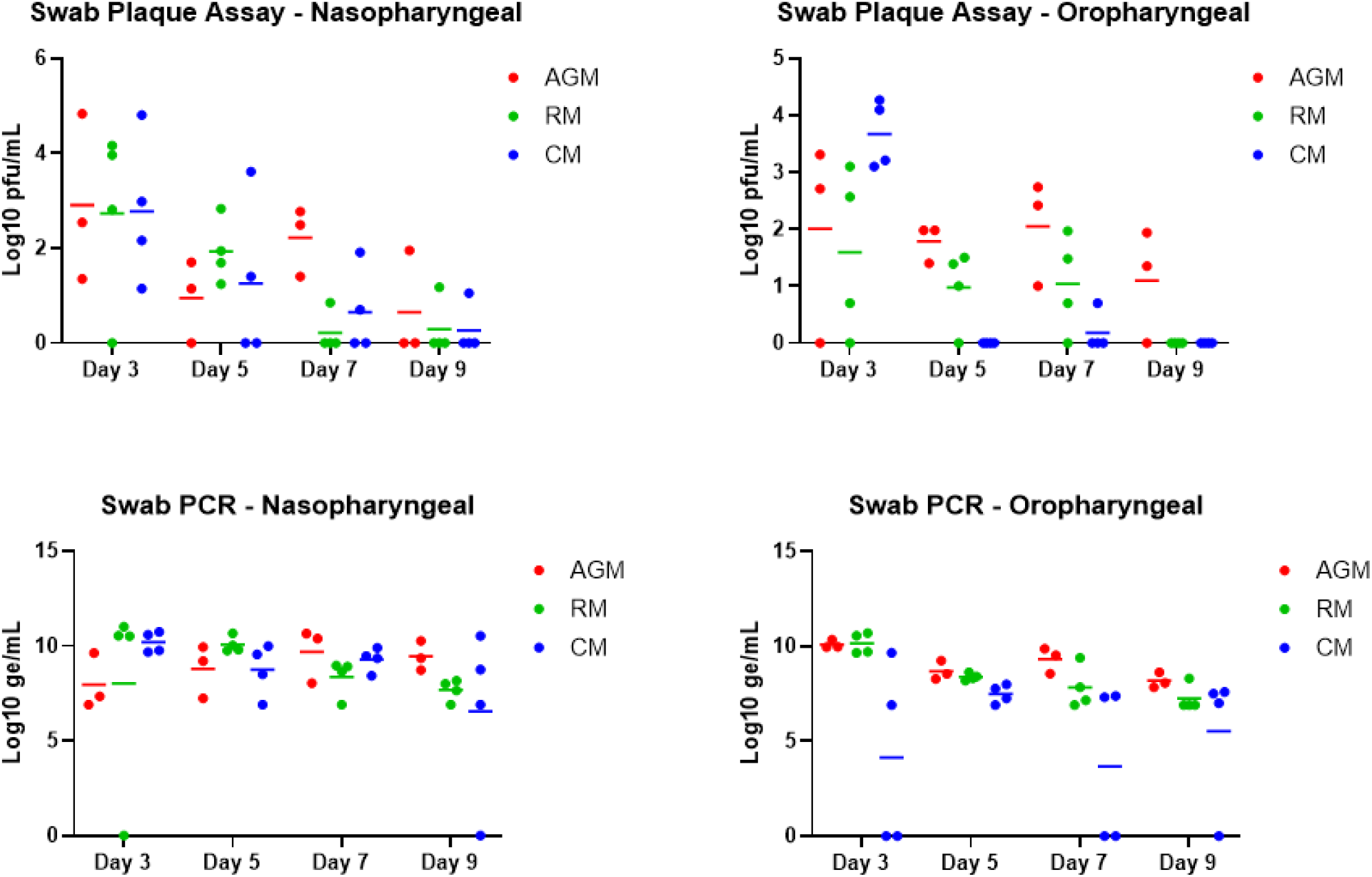
Viral RNA and live virus in oropharyngeal and nasopharyngeal swabs. SARS-CoV-2-specific qRT-PCR was performed on RNA extracted from oropharyngeal and nasopharyngeal swab clarified homogenates. Plaque assay was performed on oropharyngeal and nasopharyngeal clarified homogenates. Data are shown as Log_10_ ge/mL (qRT-PCR) or Log_10_ pfu/mL (plaque assay), with nasopharyngeal swab data for each species shown in the left panels, and oropharyngeal swab data for each species shown in the right panels.

Viability of virus in OP and NP swabs was confirmed by plaque assay (Figure 4); viable virus was not detected in any rectal swabs (data not shown). Virus was detected for most animals by Study Day 3 in both swab types, with peak levels typically detected on Study Day 3. Peak levels in NP swabs were between 2.16-4.83 Log10 pfu/mL, and peak levels in OP swabs were between 2.71-4.45 Log10 pfu/mL; significant differences between swab types and groups were not observed. For the majority of the animals, virus was no longer detected by Study Day 9; however, virus was still detected on Study Day 9 and/or 11 for a few animals (AGM 1 OP and NP, AGM 2 OP, RM 2 NP, RM 3 NP, and CM 3 NP).

Gross necropsy findings were infrequent and when they were observed, the majority of lesions were confined to the lungs. The most common finding was discoloration of one or more lung lobes for AGM and CM. In two AGM (AGMs 1 and 2), diffuse red mottling of all lobes was noted, and in CM 3 locally extensive red discoloration of all lobes was observed. CM 2 also had multifocal red to gray discoloration of the right caudal lung lobe. Two animals had adhesions within the lungs and/or thorax cavity. AGM 2 had thin, clear adhesions that extended from the left cranial to the left caudal lobe, with similar adhesions noted on the right side that involved all four lobes giving the appearance of a large, single lobe on each side. CM 4 had multiple, thin, clear adhesions that extended from the right middle lobe to the pleura of the thoracic wall.

No viral RNA was detected in any animal by *in situ* hybridization (data not shown), indicating viral infection had been cleared in these tissues at the time when animals were euthanized at the end of study.

Representative images for histology are shown in Figure 5. With the exception of CM 1, all animals had some degree of inflammation and chronic lesions (alveolar fibrosis, type II pneumocyte hyperplasia) in one or more lung lobes. The alveolar fibrosis and type II pneumocyte hyperplasia indicate previous cellular damage, either due to the viral infection and/or resulting inflammation, and these findings were more frequently noted for AGM and CM. Multinucleate cells likely attributed to viral infection were only noted for AGMs 2 and 3. Multinucleate cells possibly related to viral infection were also found in the axillary lymph node for AGM 3 and the inguinal lymph node for CM 3. Other histologic findings included the following: inflammation in the nasal septum/turbinates for AGM 2, RM 1, and CMs 2 and 3; lymphoid hyperplasia in lymph nodes, mucosa-associated lymphoid tissue, and/or gut-associated lymphoid tissue nearly all animals.

**Fig. 5.**
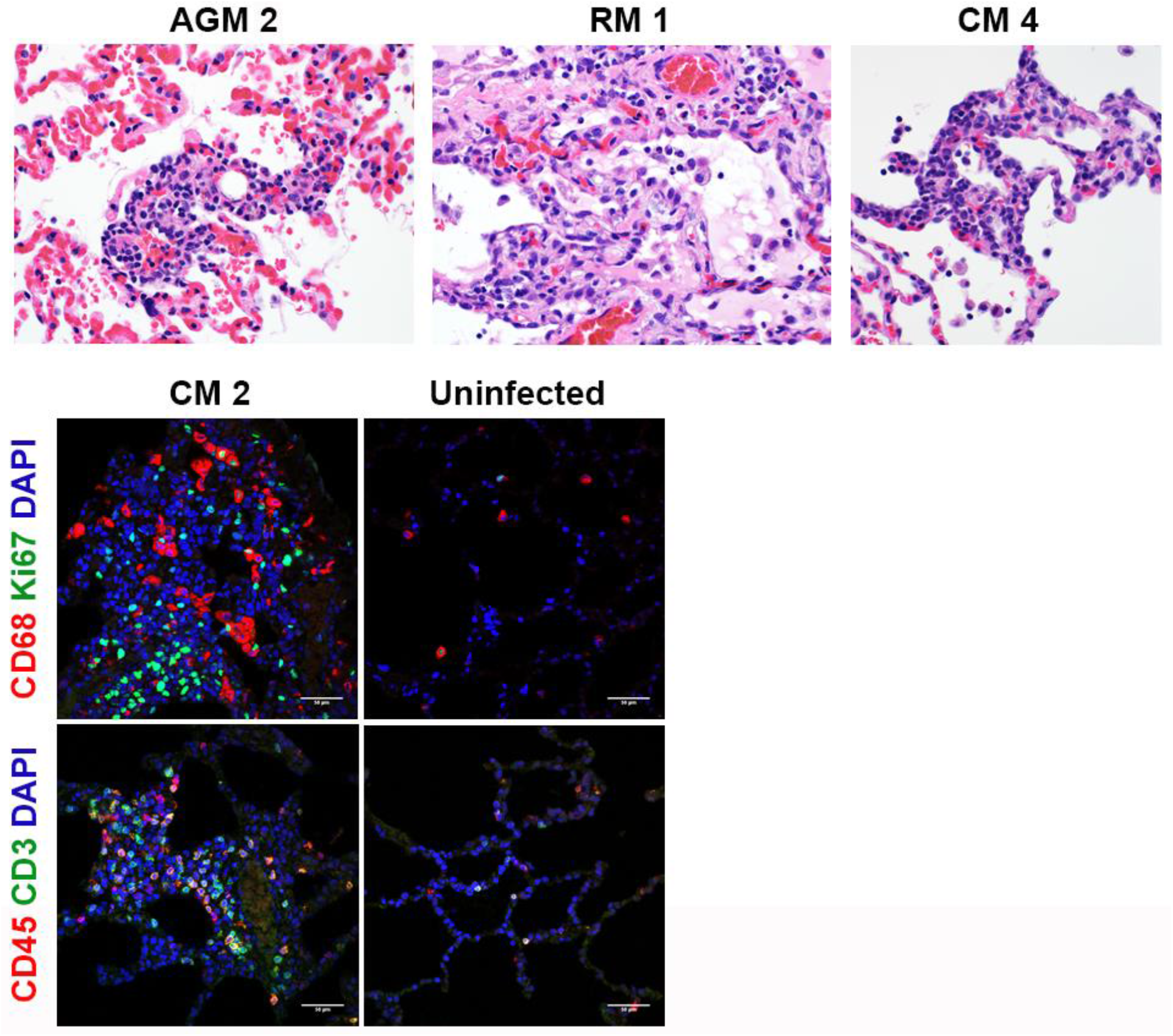
Histopathology and IFA. Necropsies, histopathology, and IFA were performed on all animals. The top panels show the following histopathology findings: AGM 2 and CM 4 = type II pneumocyte hyperplasia; RM 1 = type II pneumocyte hyperplasia and alveolar fibrosis. The bottom panels show excessive CD68^+^ macrophages (red) and Ki67^+^ proliferating cells (green), CD45^+^ leukocytes (red), and CD3^+^ T cells infiltrated in alveolar septum for CM 2 compared to uninfected control lung tissue.

Representative images for IFA are shown in Figure 5. Consistently, in comparison to historical uninfected cynomolgus macaque lung tissues, infiltration of inflammatory cells including CD68^+^ macrophages, Ki67^+^ proliferating cells, CD45^+^ leukocytes, and CD3^+^ T cells were multi-focally detected in alveolar septum of CM 2.

## Discussion

Due to the ongoing pandemic of COVID-19, it is critical that relevant animal models that recapitulate human disease characteristics are developed to facilitate immediate vaccine and treatment testing. Nonhuman primate models will be key to this testing, as data generated will be combined with human clinical data to support rapid deployment of safe and efficacious countermeasures for human use. Herein, we describe the evaluation of the first airborne SARS-CoV-2 model in macaques, and compare the results to the previously-described airborne model in AGMs (*16, 17*). The development of macaque models is critical to advance countermeasure testing efforts. Although prior studies have highlighted the potential utility of AGMs for SARS-CoV-2 testing, certain characteristics might make these nonhuman primates less than ideal candidates. AGMs are generally wild-caught and imported for use on study; the medical history for these animals is typically only available from the time of import. In addition, age can only be approximated as actual birth dates for AGMs are generally unavailable. Finally, wild-caught primates represent a diverse genetic population, resulting in inconsistent cohorts which may include animals predisposed to developing more severe disease following viral infection. For this reason, AGMs are not typically chosen as gold-standard models of viral disease. In contrast, CMs and RMs have represented critical models for countermeasure development for various pathogens over many years. This is because this population of animals is more uniform as they can be acquired from breeding colonies. Health and age information is generally available from the time of birth, decreasing the likelihood of outliers based on genetic or medical background. It is for this reason that we sought to investigate airborne exposure with SARS-CoV-2 in macaque models.

The airborne model mimics a predicted natural transmission route (*10, 18–21*), compared to other nonhuman primate models of COVID-19 which rely on direct contact inoculation of mucosal and respiratory surfaces. Exposure of AGM, RM, and CM to SARS-CoV-2 resulted in disease similar to mild cases of human COVID-19. Increased lung sounds on auscultation in association with radiographic findings consistently confirmed the presence of mild to moderate respiratory disease for all animals on study, and febrile illness was achieved for all CM.

CM developed a consistent disease, with fever being a hallmark of disease exclusive to this group. Lung infiltrates, visualized radiographically, were also common for animals in this group. The clinical disease severity score for CM was 3.00 points higher than AGM and 4.25 points higher than RM. Elevation of more than one liver-related enzyme and serum glucose were also consistently measured for CM. Overall, there were sufficient objective criteria for this group that could be easily employed to evaluate efficacy in future countermeasure efforts.

CMs have been investigated as a potential model for SARS-CoV-2, with variable disease findings noted following either IT/IN or IT/IN/ocular inoculation (*11, 22*). Importantly, fever, which represents a prominent human disease sign and which was consistently noted for CM on the present study, was not noted for any CMs on these prior studies. In addition, many of the other clinical signs of disease observed in the present study for CM were not reported for the prior studies, aside from infiltrates noted radiographically in only one of those studies. Therefore, using an airborne transmission route, we were able to generate a model of COVID-19 in CM that was more representative of human disease compared to prior work utilizing this species.

The AGM aerosol model of SARS-CoV-2 infection has been recently defined (*16, 17, 23*). In general, findings in the present study were similar to those previously described. There were two key hematologic findings that make this an attractive model, thrombocytopenia and monocytosis. Thrombocytopenia has been noted during human disease (*24*), following IT/IN inoculation (*23*), and following aerosol exposure of AGM (*17*), and this finding was consistently noted for AGM on this study. Similarly, pronounced elevation of monocytes which far exceeded levels for the other species was consistently noted for AGM. This finding has not been previously reported; the physiological relevance is unknown at this time and warrants further investigation.

Blair *et al* have been the only group thus far to describe acute respiratory distress syndrome in nonhuman primates following SARS-CoV-2 inoculation, and this was observed in only two AGM, one exposed to virus by the aerosol route and the other exposed via a combination route (conjunctival/IT/oral/IN) (*16*). It should be noted that the animals used on that particular study were “aged”. In addition, only two animals were evaluated for each exposure route, bringing into question the relevance of the finding (can severe COVID-19 actually be achieved using aged nonhuman primates, or were the two animals that developed this finding simply outliers with underlying and undetected age-related conditions that predisposed them to more severe disease).

RM have been the most extensively used SARS-CoV-2 model species, with the IT/IN or IT/IN/ocular/oral RM models being the gold standards for use in countermeasure efficacy studies (*11–13*). Clinical disease following airborne virus exposure was similar to that reported following IT/IN or IT/IN/ocular/oral inoculation. RM developed a mild respiratory disease characterized by increased respiratory noise on auscultation and changes in lung opacity radiographically. Similar to AGMs, thrombocytopenia was noted for animals in this group. Erythema around the eyes was a consistent and nearly exclusive finding for animals in this group; however, this is a relatively subjective measurement and may have been the result of ocular exposure during challenge (the eyes were not covered during airborne challenge). Rectal bleeding was another reliable finding for RM; however, this finding was most likely related to repeated swabbing, resulting in irritation of the rectal tissue, and not a SARS-CoV-2-specific finding.

Nearly all animals on the present study had some degree of inflammation and chronic lesions in one or more lung lobes at time of necropsy, and these findings were consistent with those previously reported for direct contact inoculation models (*11–13, 22, 23*). Alveolar fibrosis and type II pneumocyte hyperplasia indicative of previous cellular damage were more frequently noted for AGM and CM. By Study Day 18, most clinical disease signs (including radiographic findings) had either completely resolved or were resolving, and virus in lung tissue was below the limit of detection by ISH. Therefore, it may be important to investigate whether necropsy performed earlier in disease course could be beneficial to assessing the active disease state of primates infected by SARS-CoV-2. For countermeasure testing, where delayed disease is a possibility, the decision to shorten the length of study would obviously also need to be balanced against the risk of missing delayed-onset disease.

Elevation of serum glucose levels for AGM and CM was noted on this study, but to date this specific finding has not been described for human cases of COVID-19. It has been suggested that patients with uncontrolled diabetes are at higher risk for developing life-threatening complications following SARS-CoV-2 infection. However, there have not been any published studies comparing COVID-19 disease between diabetics and non-diabetics, and information is limited as to what role other comorbidities (high blood pressure, obesity, heart disease) may play in this observation. In short, there is no information currently available directly linking increased morbidity and mortality from COVID-19 to hyperglycemia. However, it could be predicted virus-induced elevations in glucose levels could be catastrophic for a patient already predisposed to hyperglycemia as this physiological change could lead to diabetic ketoacidosis, which alone is life threatening but in combination with other comorbidities could certainly be fatal. Considering the elevated glucose finding for AGM and CM infected with SARS-CoV-2, the role of hyperglycemia in SARS-CoV-2 pathogenesis should be further investigated.

In cases where survival cannot be used as a primary endpoint for countermeasure efficacy determinations, other statistically relevant findings are necessary, with objective criteria being preferred over subjective criteria. Herein, we describe two models (CM and AGM) for which obvious and consistent objective criteria exist that could be used for future countermeasure studies as statistical endpoints. For CM, these endpoints could be fever, the presence of lung infiltrates radiographically, and shed virus in NP and/or OP swabs. For AGM, these endpoints could be platelet levels, monocyte levels, and shed virus in NP and/or OP swabs. Although RM had fewer objective findings compared to CM and AGM, consistent subjective findings associated with shed virus in NP and/or OP swabs still make this a very valuable and utilizable COVID-19 model. Regardless of species, the present study demonstrated that macaques can be successfully infected by airborne SARS-CoV-2. Considering relevance to human disease, this study demonstrated that airborne nonhuman primate models should be strongly considered for any future countermeasure evaluations.

## Acknowledgments

The authors would like to acknowledge the members of the USAMRIID COVID-19 Protocol Working Group, the Comparative Medicine Division, and the Core Laboratory Sciences Division for their contributions during protocol conception and review. The authors would also like to thank the program managers assigned to this study for their assistance with programmatic requirements including funding execution, timeline management, and deliverable tracking.

## Author contributions

SCJ performed conceptualization, formal analysis, methodology, supervision, visualization, and writing – original draft, review, & editing. MLMP and AN performed conceptualization, funding acquisition, and writing – review & editing. AJ, JLR, XZ, JS^3^, and FR performed formal analysis, investigation, methodology, and writing – review & editing. KA and BK performed investigation, methodology, and writing – review & editing. DD, OF, JM, KB, HE, JS^2^, WS, JF, LK, SS, LW, PB, TG, CJ, JG, DN, NA, SB, FP, DD, JB, JAJ, MD, WG, and RK performed investigation and writing – review & editing.

## Data and materials availability

All data is available in the main text or the supplementary materials. A material transfer agreement would be required for the transfer of any specimens generated as part of this study.

## Supplementary Materials

Materials and Methods 5

Figure S1

Tables S1-S2

