## Supplemental Materials for "Development of a Coronavirus Disease 2019 Nonhuman Primate Model Using Airborne Exposure"

**This PDF file includes:**

Materials and Methods

Fig. S1

Tables S1 to S2

Materials and Methods

Animals

Animal research was conducted at the United States Army Medical Research Institute of Infectious Diseases (USAMRIID). Three adult *Chlorocebus aethiops* (African green monkeys) of Caribbean origin, four adult *Macaca mulatta* (rhesus macaques) of Chinese origin, and four adult *Macaca fascicularis* (cynomolgus macaques) of Chinese origin were included on this study. Genders were mixed male and female, and all animals were SARS-CoV-2 serologically naïve. All animals had passed a semi-annual physical examination and were certified as healthy by a veterinarian. Animals were acclimated in ABSL-3 animal rooms for 3 days prior to virus exposure and housed individually in 4.3 square foot cages. During the in-life portion of the study, animals were provided 2050 Monkey Chow (Harlan Teklad, Frederick, MD), fruits, and water ad libitum via an automatic watering system, and animals were given enrichment regularly as recommended by the Guide for the Care and Use of Laboratory Animals.

Ethics statement: These experiments and procedures were reviewed and approved by the United States Army Medical Research Institute for Infectious Diseases Institutional Animal Care and Use Committee (IACUC). All research was conducted in compliance with the USDA Animal Welfare Act (PHS Policy) and other federal statutes and regulations relating to animals and experiments involving animals, and adheres to the principles stated in the Guide for the Care and Use of Laboratory Animals, National Research Council, 2011. The facility is fully accredited by the Association for Assessment and Accreditation of Laboratory Animal Care, International. The animals were provided food and water ad libitum and checked at least daily according to the protocol. All efforts were made to minimize painful procedures; the attending veterinarian was consulted regarding painful procedures, and animals were anesthetized prior to phlebotomy and virus infection. Animals were humanely euthanized at the end of study by intracardiac administration of a pentobarbital-based euthanasia solution under deep anesthesia in accordance with current American Veterinary Medical Association Guidelines on Euthanasia and institute standard operating procedures.

Virus and Virus Exposure

A seed stock of SARS-CoV-2, Washington state first isolate in 2020 (WA-1/2020), designated as Lot R4717, was grown on ATCC Vero 76 cells using Stock Lot R4716 [Centers for Disease Control and Prevention (CDC) obtained]. The seed stock contains an average of 1.56×10^6^ pfu/mL of infectious virus as determined using a standardized agarose and neutral red-based assay. R4717 was fully sequenced, evaluated for sterility, tested for mycoplasma and endotoxin levels, and tested in a number of real-time reverse transcriptase polymerase chain reaction (RT-PCR) assays to include two specific for SARS-CoV-2 virus. It was determined to have no detectable mycoplasma, endotoxin or adventitious agents based on the assays and techniques used. No known contaminants were detected when sequencing the stock. Identity was confirmed by real-time RT-PCR. This stock does not contain any known contaminants and was deemed appropriate for use in testing for coronavirus therapeutics screening. On the day of exposure, Study Day 1, animals were exposed to undiluted R4717 in the USAMRIID head-only exposure system. The aerosol spray is generated using a Collison Nebulizer to produce a highly respirable aerosol (flow rate 7.5±0.1 L/minute). The system generates a target aerosol of 1 to 3 µm mass median aerodynamic diameter determined by aerodynamic particle sizer. Samples of the aerosol collected from the exposure chamber using an all-glass impinger during each exposure were assessed using a plaque assay. The exposure dose for each animal was calculated from the minute volume determined with a plexiglass whole body plethysmograph box using Buxco FinePointe software. The total volume of aerosol inhaled was determined by the exposure time required to deliver the estimated inhaled dose.

Animal Observations

Animals were evaluated cage side for signs of illness. Other observations such as biscuit/fruit consumption, condition of stool, and urine output were also documented, if possible. Observations under anesthesia (physical examinations) occurred after cage side observations on Study Days 1, 3, 5, 7, 9, 11, 15, and the day of disposition (i.e. Study Day 18). Body weights, pulse oximetry, auscultation of heart and lung sounds, radiography, blood collection, and collection of swab samples occurred during physical examinations.

Telemetry

Telemetry implants (M00; Data Sciences International, St. Paul, MN) were used to continuously monitor body temperature and activity in subject animals. Subjects were housed in individual cages in close proximity to radio frequency digital transceivers (TRX; Data Sciences International, St. Paul, MN) equipped with directional antennas pointed at the animal cages. These transceivers were connected via cat5e cables to a set of Communication Link Controllers (CLC, Data Sciences International, St. Paul, MN) to allow the digital multiplexing and the simultaneous collection of signals from all subjects. The signals were then routed over cat5e cable to data acquisition computers, which captured, reduced and stored the digital data in data files (i.e., NSS files) using the Notocord-hem Evolution software platform (Version 4.3.0.77, Notocord Inc., Newark, New Jersey). Reduced data in the NSS files was extracted into Microsoft Excel workbooks using Notocord-derived formula add-ins, and the 30 minute (min) averages were calculated for each parameter for each subject. Telemetry data collected prior to challenge was used as baseline, and provided the average and standard deviation (SD) for each 30 min daily time period of a 24 hour day.

Necropsy, Histology, *In Situ* Hybridization, and Immunofluorescence

Necropsies were conducted by a veterinary pathologist on all animals in this study. The tissue samples were trimmed, routinely processed, and embedded in paraffin. Sections of the paraffin-embedded tissues 5 µm thick were cut for histology. For histology, slides were deparaffined, stained with hematoxylin and eosin (H&E), coverslipped, and labeled. *In situ* hybridization was performed as previously described (*8*).

For immunofluorescence, the following procedures were performed. After deparaffinization and treatment with 0.1% Sudan Black B to reduce autofluorescence, tissues were heated in citrate buffer, pH 6.0 (Sigma-Aldrich, St. Louis, MO), for 15 min to reverse formaldehyde cross-links. After rinses with phosphate-buffered saline (PBS), pH 7.4 (Thermo Fisher Scientific, Waltham, MA), sections were blocked overnight with PBST (PBS+ 0.1% Tween-100) containing 5% normal goat serum (MilliporeSigma, Temecula, CA) at 4°C. Sections were then incubated with the following primary antibodies for 2 h at room temperature: rabbit polyclonal antibody against Ki67 at a dilution of 1:400 (ab15580, Abcam, Waltham, MA); rabbit polyclonal anti-CD3 antibody at a dilution of 1:200 (A045229-2, Dako Agilent Pathology Solutions, Carpinteria, CA); mouse anti-human CD68 antibody at a dilution of 1:200 (Clone KP1, Dako Agilent Pathology Solutions, Carpinteria, CA); mouse monoclonal antibody against CD45 at a dilution of 1:200 (Clone 2B11 + PD7/26, Dako Agilent Pathology Solutions, Carpinteria, CA). After rinsing in PBST, sections were incubated with secondary goat IgG Alexa Fluor 488-conjugated anti-rabbit and with goat IgG Alexa Fluor 561-conjugated anti-mouse antibody (Thermo Fisher Scientific, Waltham, MA) for 1 h at room temperature. Sections were cover-slipped using VECTASHIELD antifade mounting medium with DAPI (Vector Laboratories, Burlingame, CA). Images were captured on an LSM 880 Confocal Microscope (Zeiss, Oberkochen, Germany) and processed using open-source ImageJ software (National Institutes of Health, Bethesda, MD).

Clinical Pathology

For serum chemistries, whole blood was collected into Serum Clot Activator Greiner Vacuette tubes (Greiner Bio-One, Monroe, NC). Tubes were allowed to clot for at least 10 min and the serum separated in a centrifuge set at 1800 × g for 10 min at ambient temperature. The required volume of serum was removed for chemistry analysis using a General Chemistry 13 panel (Abaxis, Union City, CA) on a Piccolo Point-Of-Care Analyzer (Abaxis, Union City, CA). Serum was removed from the clot within 1 hour of centrifugation and was analyzed within 12 hours of collection.

For hematology, whole blood was collected into Greiner Vacuette blood tubes containing K3 EDTA as an anti-coagulant. Hematology was performed on VETSCAN® HM5 hematology analyzer (Abaxis, Union City, CA) within 4 hours of collection. In addition, 100 µL of whole blood was added to 300 µL of TRIzol® LS (Thermo Fisher Scientific, Waltham, MA) for RNA isolation for qRT-PCR.

Swab Specimen Processing

Swab samples were suspended in 1 mL of viral transport media (Hanks Balanced Salt Solution containing 2% heat-inactivated fetal bovine serum, 100 µg/mL gentamicin, and 0.5 µg/mL amphotericin B) by vortex for 15-20 seconds followed by incubation at 2-8°C for 20-25 min. Following another 15-20 second vortex, clarification was performed by centrifugation at 14,000 rpm for 30 seconds, and 100 µL of clarified supernatant was added to 300 µL of TRIzol® LS for RNA isolation for qRT-PCR. In addition, 200 µL of clarified supernatant was analyzed for infectious virus by plaque assay.

qRT-PCR

TRIzol LS whole blood and swab specimens were extracted and eluted with AVE buffer using a QIAamp® Viral RNA Mini Kit (Qiagen, Germantown, MD). The RT-PCR reaction used Invitrogen™ SuperScript® One-Step RT-PCR System with additional magnesium sulfate (MgSO_4_) added to a final concentration of 3.0 mM. Specimens were run in triplicate using a 5-µL volume. The average of the triplicates were multiplied by 200 to obtain genomic equivalents per mL, then multiplied by a dilution factor of 4 (1 part plasma to 3 parts TRIzol LS) for the final reported value. The genomic equivalents were determined using a standard curve of synthetic RNA of known concentration. Sequences of primers and probes used are as follows:

| Forward primer sequence (5’-3’) | TTACAAACATTGGCCGCAAA |
| --- | --- |
| Reverse primer sequence (5’-3’) | GCGCGACATTCCGAAGAA |
| Probe sequence (5’-3’) | ACAATTTGCCCCCAGCGCTTCAG |

Plaque Assay

Challenge dose and infectious virus in swab samples were determined by agarose plaque assay, run on fresh (i.e. not frozen then thawed) samples the day of collection. Required dilutions of each specimen were prepared in virus diluent [1X minimum essential media (Corning Life Sciences, Pittston, PA) containing 10% fetal bovine serum (Cytiva Life Sciences, Marlborough, MA), 1% GlutaMAX (Gibco, Waltham, MA), and 1% NEAA (Sigma, St. Louis, MO)] in duplicate (swab specimens) or triplicate (all-glass impinger specimens from Study Day 1), were added to plates containing Vero 76 cells (ATCC, Manassas, VA) at ≥85% confluency. Two days later, the cells were stained with neutral red (Thermo Fisher Scientific, Waltham, MA), and plaque counts were obtained the day after staining.

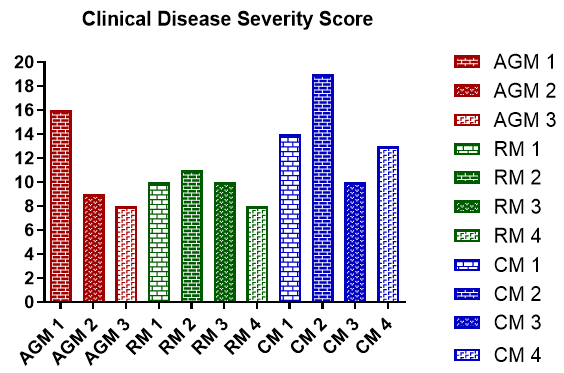

Fig. S1. Clinical disease severity. Disease severity based on clinical signs was graded using a scoring system that can be found in Table S1.

| **Parameter** | **Description** | **Score** |
| --- | --- | --- |
| Responsiveness | Alert, responsive, normal species specific behavior | 0 |
|  | Slightly diminished general activity, subdued but responds normally to external stimuli | 1 |
|  | Withdrawn, may have head down, upright fetal posture, hunched, reduced response to external stimuli | 2 |
|  | Prostrate but able to rise if stimulated, or dramatically reduced response to external stimuli | 6 |
|  | Persistently prostrate (unable to rise when stimulated), or severely or completely unresponsive, | 8 |
| Discharge | Nasal or ocular | 2 |
|  | Nasal and ocular | 4 |
| Integument | Rash or unusual bruising | 2 |
|  | Rash and unusual bruising | 4 |
| Respiratory Function | Normal - no apparent changes in breathing, 30-50 breaths per minute, and no cough | 0 |
|  | Mild dysfunction - 50-65 breaths per minute, increased respiratory sounds on auscultation, or isolated cough | 2 |
|  | Moderate dysfunction - increased effort of breathing (abdominal breathing and/or nasal flare), 66-80 breaths per minute, or apparent cough | 4 |
|  | Severe dysfunction – continuous gasping, open mouth breathing + abdominal breathing, and/or cyanosis | 8 |
| Food consumption | Evidence of biscuit and fruit consumption | 0 |
|  | No evidence of biscuit or fruit consumption | 1 |
|  | No evidence of biscuit and fruit consumption | 4 |
| Stool/GI | Normal | 0 |
|  | Soft or liquid stool present, palpable gas or palpable fluid in GI region | 1 |
|  | Rectal bleeding | 2 |
| Body Temperature | Normal | 0 |
|  | Temperature elevated (>3 standard deviations above baseline) | 2 |
|  | Fever (>1.5°C above baseline) | 4 |

| **Parameter** | **Description** | **Score** |
| --- | --- | --- |
| Heart rate | Normal (up to 19 BPM over baseline) | 0 |
|  | Mild tachycardia (20-39 BPM over baseline) | 1 |
|  | Moderate tachycardia (40-69 BPM over baseline) | 2 |
|  | Severe tachycardia (>70 BPM over baseline) | 3 |
| Respiratory rate | Normal - 30-50 breaths per minute | 0 |
|  | Mild tachypnea- 50-65 breaths per minute | 2 |
|  | Moderate tachypnea - 66-80 breaths per minute | 4 |
|  | Severe tachypnea - >80 breaths per minute | 6 |
| SpO_2_ | Normal (95-100%) | 0 |
|  | Mildly decreased (90-94%) | 2 |
|  | Moderately decreased (87-89%) | 4 |
|  | Severely decreased (<87%) | 6 |
| Body weight | Normal (0-3% loss) | 0 |
|  | Mild (4-9% loss) | 1 |
|  | Moderate (10-16% loss) | 2 |
|  | Severe (>16% loss) | 3 |
| Clotting following blood collection | Normal – site of venipuncture clots quickly | 0 |
|  | Abnormal – noticeable increase in time required for the animal to clot following venipuncture | 2 |
| Gait, locomotion, and balance | Normal | 0 |
|  | Mild – shaking | 1 |
|  | Moderate – difficulty grasping items, or difficulty moving around cage | 4 |
|  | Severe – unable to moving around cage, unable to climb, or falling down | 6 |
| Lymphadenopathy | Absent | 0 |
|  | Present | 1 |

| **Parameter** | **Description** | **Score** |
| --- | --- | --- |
| Conjunctival Erythema | Absent | 0 |
|  | Present | 1 |
| Radiographic Findings | Normal | 0 |
|  | Mild – opacity/glassy appearance or infiltrates in at least one lung lobe | 1 |
|  | Moderate – opacity/glassy appearance or infiltrates in at least two lung lobes | 2 |
|  | Severe – opacity/glassy appearance or infiltrates in at least three lung lobes | 3 |

Table S1. Nonhuman Primate COVID-19 Clinical Disease Severity

| Animal Number | AGM 1 | AGM 2 | AGM 3 | RM 1 | RM 2 | RM 3 | RM 4 | CM 1 | CM 2 | CM 3 | CM 4 |
| --- | --- | --- | --- | --- | --- | --- | --- | --- | --- | --- | --- |
| WBC  (+/-) | X  (-39%)  (+30%) | X  (-37%) | X (+69%) | X  (-28%) | X  (+36%) | X  (-29%)  (+31%) | X (+28) | X  (-29%) | X  (-44%) | X  (-52%) | X  (-29%) |
| NEU  (+/-) | X  (-58%) | X  (-56%) | X  (-55%)  (+80%) | X  (-30%) | X  (+44%)  (-34%) | X  (-58%)  (+58%) | X  (-46%) | X  (-31%) | X  (-59%) | X  (-60%) | X  (-34%) |
| LYM  (+/-) | X  (-31%)  (+52%) | X  (-99%)  (+52%) | X  (-95%)  (+155%) | X  (-27%) | X  (-29%)  (+33%) | X  (-68%)  (+32%) | X  (-41%)  (+79%) | X  (-54%)  (+96%) | X  (-43%)  (+108%) | X  (+90%) | X  (-72%) |
| MON  (+/-) | X  (-33%)  (+167%) | X  (+5,633%) | X  (+4,300%) | X  (+80%) | X  (+167%) | X  (+67%)  (-50%) | X  (+125%)  (-25%) | X  (+220%) | X  (+314%) | X  (-50%)  (+33%) | X  (-55%)  (+64%) |
| PLT  (+/-) | X  (-51%) | X  (-43%) | X  (-52%) | X  (-42%) |  |  |  |  |  |  | X  (-38%) |
| ALT  (+) | X  (+84%) | X  (+36%) |  | X  (+64%) |  | X  (+42%) | X  (+48%) | X  (+76%) | X  (+85%) | X  (+34%) | X  (+48%) |
| ALB  (-) |  |  |  |  |  |  |  |  | X  (-39%) |  |  |
| ALP  (+) | X  (+32%) |  | X  (+37%) |  |  |  |  | X  (+165%) |  |  | X  (+35%) |
| AST  (+) | X  (+127%) | X  (+117%) | X  (+29%) | X  (+29%) | X  (+56%) |  | X  (+49%) | X  (+153%) | X  (+46%) | X  (+79%) | X  (+126%) |
| GGT  (+) | X  (+40%) | X  (+30%) | X  (+57%) |  |  |  |  | X  (+129%) |  |  |  |
| GLU  (+/-) | X  (+63%) | X  (+69%) | X  (+41%) |  |  |  |  |  | X  (+33%) | X (+26%) | X  (+45%) |

X = >25% change from baseline for a given parameter for at least 1 time point

() = maximum percent change from baseline noted for an animal

**Table S2. Summary Clinical Pathology Findings**
